# Direct DNA sequencing of ‘*Candidatus* Liberibacter asiaticus’ from *Diaphorina citri*, the Asian citrus psyllid, and its implications for citrus greening disease management

**DOI:** 10.1101/2022.01.28.478250

**Authors:** Steven Adam Higgins, Marina Mann, Michelle Heck

## Abstract

The Asian citrus psyllid, *Diaphorina citri* is an invasive insect ^1^ and a vector of ’*Candidatus* Liberibacter asiaticus’ (*C*Las), a bacterium whose growth in *Citrus* species results in citrus greening disease ^2,3^. Methods to enrich and sequence *C*Las from *D. citri* often rely on biased genome amplification ^4^ and nevertheless contain significant quantities of host DNA ^5,6^. To overcome these hurdles, we developed a simple <u>p</u>re-treatment <u>D</u>Nase and filtration (hereafter PDF) protocol to directly sequence *C*Las and the complete, primarily uncultivable, microbiome from *D. citri* adults. The PDF protocol yielded *C*Las abundances upwards of 60% and enabled detection of 156 genetic variants in these strains compared to progenitor strains in Florida, which included prophage encoding regions with key functions in *C*Las pathogenesis, putative antibiotic resistance loci, and a single secreted effector. These variants suggest laboratory propagation of *C*Las may result in different phenotypic trajectories among laboratories, and may confound *C*Las physiology or therapeutic design and evaluation if these differences remain undocumented. Finally, we obtained genetic signatures affiliated with *Citrus* nuclear and organellar genomes, entomopathogenic fungal mitochondria, and commensal bacteria from laboratory-reared and field-collected *D. citri* adults. Hence, the PDF protocol can inform agricultural management strategies related to pathogen evolution ^7^, insect microbiome surveillance ^8^, antibiotic resistance screening ^9^, and gut content analysis ^10^.

## Introduction

The abundance of host DNA in nucleic acid extracts is a significant barrier to cost-effective sequence analysis of host-associated microbiomes ^11^. In addition, the non-uniform abundance and spatial organization of a host’s microbial populations significantly impedes direct sequence analysis of host-associated microbiomes. Sequencing effort is therefore directed towards host DNA and limits the cost-effectiveness and utility of shotgun sequence data in microbiome analysis ^12^. To help overcome these issues, a combination of pre- and post-DNA extraction methods have been devised to limit host DNA in nucleic acid extracts ^11^. These methods include differential lysis of host tissues followed by exonuclease treatment ^11^, host DNA depletion via methylation status followed by multiple displacement amplification ^13–15^, use of targeted hybridization probes for selective enrichment of microbial DNA ^16,17^, and cell enrichment by cultivation in alternative hosts ^18^. While useful, these methods are relatively costly, laborious, and can restrict analysis to a single taxon (e.g., probe hybridization) or introduce biases (e.g., using multiple displacement amplification) that limit the types of downstream analyses afforded by shotgun sequence data ^4^.

One host-microbe system that illustrates the impediment of host DNA in microbiome analyses is that of citrus greening disease pathosystem ^6,19^. Citrus trees worldwide are threatened by this rapidly spreading disease, which over a period of years significantly reduces citrus fruit yield, marketability, and eventually results in tree decline and death ^2,3,20^. Citrus greening disease is estimated to cost some U.S. citrus growers $1 billion annually and has reduced Florida citrus production by over 70% since the early 2000s ^21,22^. In the US, the disease is attributed to the growth of the obligately biotrophic, gram-negative bacterium ‘*Candidatus* Liberibacter asiaticus’ (*C*Las) within citrus trees, which induces phloem-plugging and results in the characteristic citrus greening pathology of aborted ripening, blotchy leaf mottle, and tree root decline ^2,20,22^. *C*Las is proposed to be transmitted by its insect vector, *Diaphorina citri* Kuwayama (Hemiptera: Liviidae) the Asian citrus psyllid in a circulative propagative manner which involves extensive interaction with and replication the vector’s tissues ^23,24^. *D. citri* acquire *C*Las during the nymphal stage and become competent to transmit as adults when *C*Las titer reaches high levels in the insect’s salivary glands ^24^. Like the majority of phloem-feeding hemipteran insects, *D. citri* harbor three known bacterial symbionts whose functions in their biology have been inferred from genome sequencing: *Wolbachia pipientis* strain wDi ^25^, ‘*Candidatus* Profftella armatura*’* a symbiont with predicted nutritional and defensive functions ^26,27^ and ‘*Candidatus* Carsonella rudii’ a nutritional symbiont ^25,28^. These latter two symbionts are essential for *D. citri* survival but their role in *C*Las transmission is speculative ^29^. *D. citri* have spread *C*Las throughout citrus growing regions in the U.S. since its initial discovery in Florida in 2005 ^2^. While we report on the impacts of citrus greening to the US citrus industry for simplicity, we would like to emphasize that this disease presents a significant threat to citrus production globally ^3,30^. Particularly alarming is that a handful of countries globally produce a majority of the world’s citrus crops ^3,30^, and that some at-risk countries possess large swaths of habitat favoring the spread of citrus greening disease ^30,31^.

Historically, the cultivation of microbial plant pathogens has facilitated their experimentation and comparative sequence analysis, or design of therapeutics to control their spread, while currently uncultivable *Ca*. Liberibacter pathogens (like *C*Las) are difficult to study due to their obligately biotrophic lifestyle ^32^. Since attempts to isolate and continuously culture *C*Las have been unsuccessful ^33^, high-throughput sequencing has become an invaluable tool for hypothesis generation and performing metabolic, comparative genomics, and evolutionary analyses of this pathogenic bacterium ^2,34,35^. Nevertheless, many hurdles to efficient and direct sequencing of *C*Las populations from both citrus and *D. citri* exist ^6,19^. For example, due to the overabundance of host DNA in nucleic acid extracts from citrus and psyllid samples, multiple enrichment strategies that include deep sequencing, multiple displacement amplification of host-depleted DNA, and antibody- or RNA-probe capture hybridization are employed to facilitate sequence analysis of *C*Las and other *Ca.* Liberibacter populations ^16,19,33,36,37^.

Robust and facile sequencing of *C*Las and other pathogenic, host-associated microbial populations is necessary to keep pace with the rapid development and application of therapeutics to prevent their spread ^38^. While a variety of antimicrobials have been applied to prevent the spread of citrus greening disease ^21,38^, we are currently unable to perform cost-effective and rapid sequencing of *C*Las populations to screen for genetic resistances to therapeutics if and when they arise. Furthermore, methods to identify and compare *C*Las populations have been limited to a few dozen samples that have been sequenced worldwide ^34^. This small sample size limits the identification and development of targeted therapeutics and hampers our ability to contrast standing genetic variation detected in laboratory-propagated *C*Las populations with those actively spreading in agricultural settings. Thus, we developed a simple and efficient pre-treatment DNase and filtration (PDF) protocol to sequence *C*Las and other bacterial populations directly from individual members of their insect host, *D. citri*. We compare and contrast our method with existing sequencing methods used to sequence *C*Las populations and present results of our genomic analyses enabled by the PDF protocol on both lab-reared and field-collected *D. citri* sampled from a minimally-managed citrus grove in Florida. Specifically, we report on replication rate estimates and genetic polymorphisms detected among microbial populations within individual *D. citri* adults and discuss their implications for citrus greening disease management and insect microbiome research. The PDF protocol enables simultaneous analysis of all microbial community members in a single *D. citri* adult, from symbionts to commensals, that will greatly inform diverse research fields, such as therapeutics development and application ^9^, holobiont theory ^39^, and ecological and evolutionary investigations of *D. citri* and other agriculturally relevant pests ^7,8^.

## Methods

### Rearing and collection of laboratory-reared *D. citri* adults

Colonies of the Asian citrus psyllid, *Diaphorina citri* Kuwayama (Hemiptera: Liviidae), were reared in controlled growth chambers on *C*Las-infected *Citrus medica* (citron) plants under 14:10 h light:dark cycle at 28□. Citrus plants were regularly monitored by lab technicians and maintained by pruning and watering on a weekly basis or as needed. Individual *D. citri* adults from *C*Las-infected citron colonies were bulk-collected using a sterile 25 ml serological pipette under vacuum into a sterile 50 ml falcon tube. The falcon tube containing *D. citri* adults was then transferred onto ice to immobilize the insects prior to handling and sorting. Immobilized *D. citri* were then transferred into a plastic weigh boat on ice and individual *D. citri* were placed into sterile 2 ml microcentrifuge tubes using a 70% (v/v) ethanol-wiped, fine-tipped paintbrush and kept on ice until processing for DNA extraction.

### Field collection of *D. citri* adults

With permission from a local citrus grower, *D. citri* adults were collected from *Citrus sinensis* (Valencia orange) trees in a minimally managed citrus grove located in St. Lucie county, FL, in early October of 2018. This grove did not utilize pesticides and natural enemies of *D. citri* were frequently observed (e.g., spiders and parasitoid wasps) ^40^. Individual *D. citri* were collected from citrus leaf surfaces using mouth aspiration into sterile borosilicate glass collection vials from a few dozen citrus trees. The individual *D. citri* were then transferred into 95% ethanol (v/v) and kept stored on ice. In the lab, the ethanol was decanted and the *D. citri* adults were air dried on Kimwipes until storage at -80 °C prior to DNA extraction.

### Pre-treatment DNase and filtration (PDF) protocol and sequencing

To each tube containing an individual *D. citri* adult, we added 100 µl of master mix containing 8 U DNase I (TURBO DNA-free kit, catalog no. AM1907, Thermo Fisher Scientific, Waltham, MA, USA), 1X DNase I buffer, 2X ProteoGuard EDTA-free protease inhibitor cocktail (catalog no. 635673, Takara Bio USA, Inc., San Jose, CA, USA), and DNase and RNase-free RPI molecular grade water (catalog no. 248700-500.0, Neta Scientific, Inc., Hainesport, NJ, USA). Psyllids were then manually homogenized within each tube for 30 sec using an autoclaved Bel-Art polypropylene microcentrifuge pestle (catalog no. 199230001, SP Industries, Inc., South Wayne, NJ, USA). Following homogenization, samples were briefly centrifuged and gently flicked with a finger to mix the tube contents, followed by incubation in a water bath at 32□ for 2 h. Tubes were gently mixed by hand agitation during incubation at 30 min intervals. Following this incubation, samples were placed on ice and 100 µl of sterile 1X TE buffer (10 mM Tris HCl, 1 mM EDTA, pH 8.0) was added. The total mixture was then transferred to autoclaved, 1.5 ml plastic microcentrifuge tubes containing a 5 µm pore size PVDF filter cartridge (catalog no. UFC30SV00, MilliporeSigma, Burlington, MA, USA) followed by centrifugation at 8,000 x *g* for 3 min. The supernatant was carefully removed by pipetting and the resulting cell pellet was placed on ice. DNA was then extracted from the cell pellet by addition of 600 µl of buffer RLT (catalog no. 79216, Qiagen Sciences, Inc., Germantown, MD, USA) followed by gentle pipetting to reconstitute the cell pellet. The mixture was then transferred to a sterile 2 ml microcentrifuge tube containing pre-autoclaved and air-dried 0.1 mm zirconia-silicate beads (catalog no. 11079101Z, Bio Spec Products, Inc., Bartlesville, OK, USA) and homogenized using a mixer mill 400 (Retsch GmbH, Haan, Germany) at 25 Hz for 1 min. A 450 µl aliquot of the homogenate was then transferred to a sterile 2 ml EconoSpin mini spin column (catalog no. 1910-250, Epoch Life Science, Inc., Sugar Land, TX, USA) containing 450 µl of 70% (v/v) molecular grade ethanol (catalog no. 3916EA, Decon Laboratories, Inc., King of Prussia, PA, USA) and gently pipetted up and down to mix. The samples were then centrifuged at 8,000 rpm for 1 min, the flow-through discarded, and an additional 700 µl of 75% (v/v) molecular grade ethanol was added to the column followed by centrifugation at 8,000 rpm for 30 sec and the flow-through discarded. This process was repeated an additional two times, followed by a 2 min centrifugation of the empty column at 12,000 rpm to remove residual ethanol. The DNA was eluted off the column using 25 µl of 0.1X TE buffer and DNA concentration and quality evaluated using a NanoDrop 2000 instrument (Thermo Fisher Scientific, Inc.). DNA samples were provided to the <u>Microbial Genome Sequencing Center</u> (Pittsburgh, PA, USA) for 2 x 151 bp Illumina sequencing at 200 Mbp or 1 Gbp depth offerings. Of note, the DNA concentrations range of these samples were very low (∼0.1 – 0.6 ng/µl). In earlier iterations of this workflow, preparation of sequencing libraries were unsuccessful using DNA samples not stored or shipped on ice. Any DNA derived from this method should therefore be kept on ice when handled for in-house sequencing or shipped on ice or dry ice to sequencing facilities to ensure stability of the minute DNA quantities present in the processed samples. In addition to the PDF protocol, *D. citri* adults were placed into a sterile 2 ml microcentrifuge tube containing two sterile 3.2 mm steel beads and were snap frozen in liquid nitrogen. Next, samples were cryoground on the Retsch mixer mill 400, their DNA extracted, and samples submitted for sequencing as described above. We refer to this last step as the “Direct protocol” and compare and contrast results from both the PDF and Direct protocols in the results section (Supplementary Fig. 1).

### Quantitative Polymerase Chain Reaction (qPCR) analysis

Prior to sequencing, DNA samples were analyzed by qPCR using gene specific PCR primers to monitor *C*Las enrichment relative to DNA from its *D. citri* host. The PCR primers targeted either the *C*Las 16S rRNA gene (5’-TCGAGCGCGTATGCGAATAC-3’ and 5’-GCGTTATCCCGTAGAAAAAGGTAG -3’ forward and reverse primers, respectively) or the *D. citri* wingless (*wg*) gene (5’-GCTCTCAAAGATCGGTTTGACGG -3’ and 5’-GCTGCCACGAACGTTACCTTC-3’ forward and reverse primers, respectively). PCR amplification was performed using 2X PowerSYBR green master mix (catalog no. 4367695, Thermo Fisher Scientific, Inc.). Briefly, the qPCR reaction was composed of 5 µl of 2X PowerSYBR green master mix, 0.25 µl each of the forward and reverse primers (10 µM stock), 0.125 µl of molecular grade bovine serum albumin (20,000 ng/µl stock, catalog no. B9000S, New England Biolabs, Inc., Ipswich, MA, USA), 1 µl of DNA, and 3.375 µl of molecular grade water. The qPCR amplification was performed on an Applied Biosystems QuantStudio 6 Flex Real-Time PCR System (Thermo Fisher Scientific, Inc.). Thermocycler conditions consisted of 2 min incubation at 50 °C followed by 10 min incubation at 95 °C and 40 cycles at 95 °C for 15 s and 55 °C for 1 min followed by denaturation at 95□ for 15 sec and melt curve analysis from 60□ (1 min) to 95□ (15 sec) at a rate 0.05 □/sec. The qPCR Cq values were converted to *C*Las 16S rDNA gene copies using duplicate standard curves comprised of eight, ten-fold serial dilutions of a known concentration of pUC57 plasmid (2,710 bp length) containing a *C*Las 16S rDNA gene fragment (427 bp length) prior to comparison with *D. citri* wingless genes. A similar methodology was used to calculate *D. citri wg* gene copies, except that genomic DNA was extracted as previously described ^41^ from a pool of *D. citri* adults (n = 10) followed by six ten-fold serial dilutions to construct an *D. citri* genomic DNA standard curve. An estimated *D. citri* genome size of 485,705,802 bp was used to calculate *wg* gene copies for a given quantity of DNA in the dilution series. The range of *C*Las 16S rDNA gene copies in the standard curve dilution series were 8.4 x 10^2^ to 8.4 x 10^9^, and the assay possessed a slope of -3.13, y-intercept of 39.70, R^2^ of 0.986, and efficiency of 108.7%. The range of *wg* gene copies in the standard curve dilution series were 2.5 x 10^-1^ to 2.5 x 10^5^, and the *wg* qPCR assay possessed a slope of -2.98, y-intercept of 37.96, adjusted R^2^ of 0.95, and efficiency of 118.37%.

### Data acquisition and analysis

Raw read data sets of *C*Las genomes derived from citrus tissues were accessed from the National Center for Biotechnology and Information (NCBI) Sequence Read Archive (SRA) ^42^ using accession numbers SRR016816, SRR7050882, SRR8177702, SRR9641236, and SRR9673120. These Illumina reads represent *C*Las genomic data from an unknown *C*Las strain identified in *Citrus sinensis* in Florida, USA, strain TX1712 collected from *C. sinensis* in Texas, USA ^43^, strain YNJS7C identified in *C. sinensis* from Yunnan province, China ^44^, strain AHCA17 identified in *C. maxima* (pummelo) in California, USA ^45^, and strain TaiYZ2 collected from *C. maxima* (pummelo) in Songkhla province, Thailand ^46^, respectively.

The latest *D. citri* (version 3) and *C. medica* (version 1, HZAU) and *C. sinensis* genome (version 2, HZAU) assemblies were downloaded from <u>citrusgreening.org</u> and <u>citrusgenomedb.org</u>, respectively. All raw fastq sequence files were processed with FaQCs (v2.10) ^47^ to trim (phred score cutoff ≥ 30) and remove low quality and short (<50 bp length) Illumina reads. Individual samples and a total combined sample set were assembled or co-assembled, respectively, using the SPAdes assembler (v3.14.0) ^48^ with k-mer sizes of 33, 55, and 77 for individual samples and 77, 99, and 127 for the co-assembly (all raw reads from individual samples pooled prior to assembly). Trimmed Illumina reads were aligned to assembled scaffolds using bowtie2 (v2.3.4) ^49^ and coverage calculated using samtools (v1.9) ^50^ mpileup and coverage modules with default settings. All scaffolds were aligned to the nt database using BLAST+ (v2.11.0) ^51^ and taxonomic assignments inferred using BASTA (v 1.4.1) ^52^ “sequence” module with settings -i 80 -b True -p 90. Blobtools (v1.1.1) ^53^ was used to visualize GC%, coverage, and taxonomic assignments of assembled scaffolds (data not shown), and inspired the plots used in portions of the results and Supplemental Information. Automatic binning of assembled scaffolds into coherent operational taxonomic units was performed using LAST (version 1145) ^54^ alignments and BASTA taxonomic assignments, followed by bin consistency analysis using the CheckM (v1.1.3) tool’s “lineage_wf” module to assess the bin completeness and consistency metrics ^55^. QUAST (v5.1.0rc1, settings -m 100 -f –rna-finding -b) ^56^ was used to compare assembled genome bins of *D. citri* symbionts with high quality, complete reference symbiont genomes and report assembly quality statistics.

The iRep tool ^57^ with default settings was used to estimate the index of replication (average, instantaneous population replication rate) against representative reference genomes for *C*Las, *Wolbachia pipientis* strain wDi, ‘*Candidatus* Profftella armatura’ (hereafter Prof), and ‘*Candidatus* Carsonella ruddii’ (hereafter Car) populations detected within individual *D. citri* adults processed in the present study. Snippy (v4.6.0) ^58^ was used with parameters –mincov 10 – minfrac 9 and –minqual 100 to detect, enumerate, and compare genetic polymorphisms among *D. citri* symbionts and reference symbiont genomes. Gubbins (v2.4.1) ^59^ was used with settings --tree_builder raxml --raxml_model GTRGAMMA and additional default settings to flag and generate a recombination-free single nucleotide polymorphism (SNP) alignment among whole *C*Las genomes assembled in the present study and publicly available reference genomes, which was supplied to IQ-TREE (v1.6.1) ^60^. IQ-TREE was run with settings -m K3Pu+F -bb 1000 - bnni to generate a SNP phylogeny. Some *C*Las genomes (both publicly available and in the present study) were highly fragmented and/or missing >25% of the genome and were not included in the phylogenetic analysis of SNPs. Statistics were performed in R 4.1.2 ^61^. Vizualization of SNP data and phylogeny was performed using the R packages BioCircos v0.3.4^62^ and ggtree ^63^.

### Data availability

Raw Illumina sequence data were deposited to NCBI Sequence Read Archive under BioProject no. PRJNA779156. The assembled data, their taxonomic assignments, SNP alignment and phylogeny, and all data inputs required to reproduce the figures presented in the manuscript are accessible in the figshare repository https://doi.org/10.6084/m9.figshare.c.5810090.v1.

### Code availability

All scripts used to process data and generate Fig.s can be found in the figshare repository https://doi.org/10.6084/m9.figshare.c.5810090.v1.

## Results

We introduced a variety of methods, including deep sequencing, multiple displacement amplification, methylation-based DNA depletion, and probe capture hybridization, that are available to enrich and sequence *C*Las from both citrus and *D. citri* samples. While limited comparisons using reported total read counts or the number of read alignments from the literature were possible (Supplementary Table 1, Fig. 1a), access to raw sequence data (i.e., Illumina reads) for most published *C*Las genomes were unavailable on public sequence repositories, such as the SRA ^42^. Therefore, we compared sequence data produced using the PDF protocol, which uses pre-extraction DNase treatment and filtration to remove host DNA, with data produced by direct DNA extraction from *C*Las-infected *D. citri* adults (Direct protocol) (Supplementary Fig. 1). We reasoned that robust comparisons of *C*Las-infected *D. citri* adults with variable *C*Las titers could be made using the PDF and Direct protocols whereas the same analysis was not feasible with other published methods since publicly available raw data sets were lacking. Regardless, improvements on baseline host contamination facilitated by the PDF protocol were made possible by comparison to the Direct protocol.

**Fig. 1.**
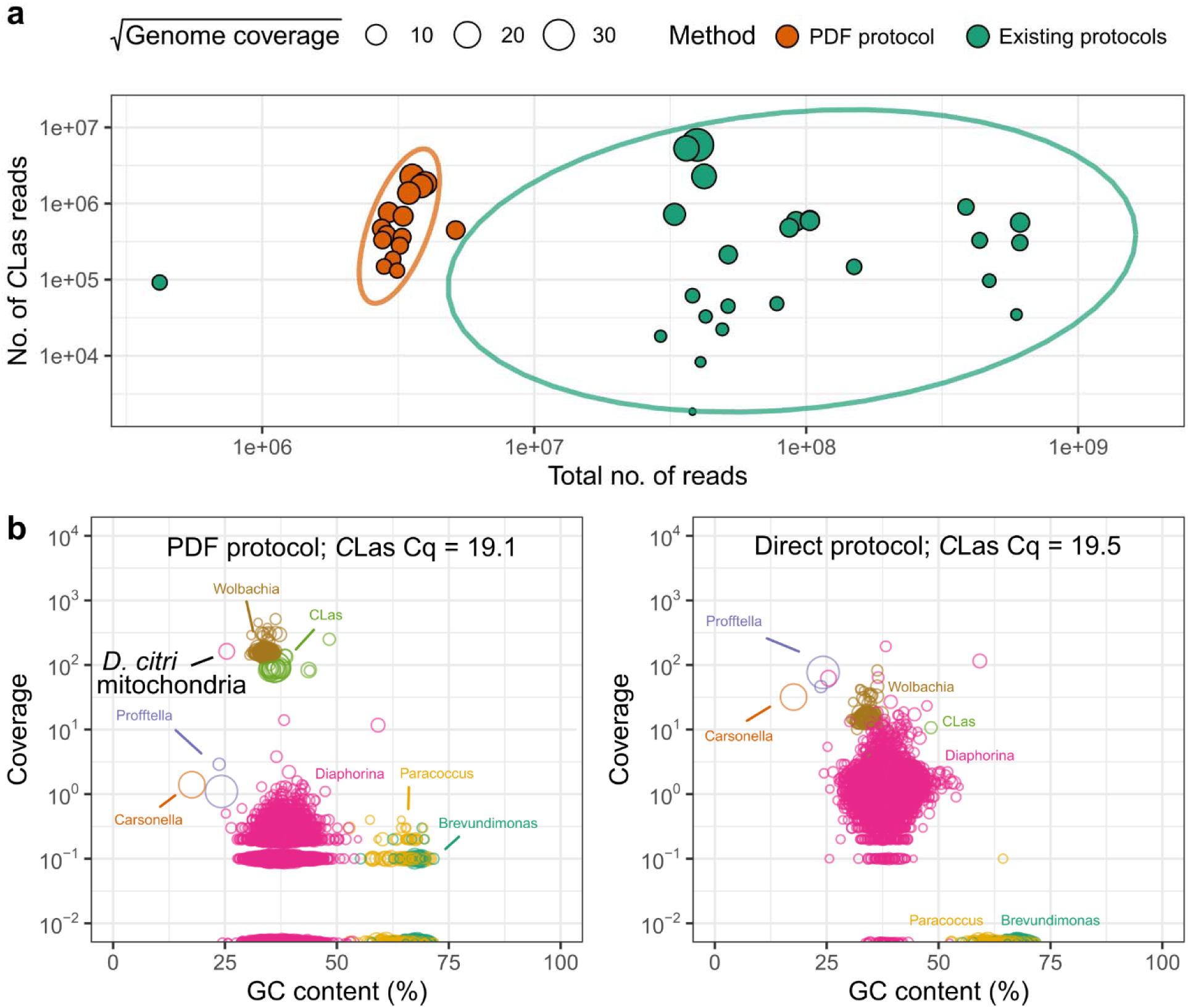
Comparison of *C*Las and additional microbial community member abundances between (a) the PDF protocol and existing *C*Las sequencing protocols in the literature or (b) between the PDF and Direct protocols using laboratory-reared *Diaphorina citri* adults with equivalent *C*Las titers. In the top panel, the PDF protocol (orange circles) generates equivalent or greater amounts of *C*Las data with orders of magnitude less sequencing data than existing protocols (green circles), reducing sequencing costs and increasing useable microbiome data. (b) In addition to increasing *C*Las abundance relative to the host, the PDF protocol also improves detection and analysis of *Wolbachia pipientis* strain wDi and other commensal bacteria (*Paracoccus* sp. and *Brevundimonas* sp.). For *D. citri* individuals with comparable *C*Las titers (assessed via qPCR; text inside bottom panels), the relative abundance of *C*Las and *Wolbachia* in these samples is an order of magnitude greater with the PDF protocol compared to directly sequencing DNA extracted from an individual *D. citri* adult (“Direct protocol” in bottom right panel). The size and color of points in (b) represent the length and taxonomic affiliations of contigs generated from the coassembly of sequence reads from all lab-reared *D. citri* individuals processed in the present study. Hence, taxa with fewer, larger circles represent larger, more contiguous assemblies. Plots in (b) were inspired by BlobTools ^53^.

To determine the effect of the PDF protocol on *C*Las titers relative to the insect host *D. citri* DNA, we quantified *C*Las 16S rRNA and *D. citri* wingless (*wg*) gene copies in DNA samples generated using the PDF protocol and compared these values with the Direct protocol. The values of *C*Las 16S rRNA gene copies for the PDF and Direct protocols ranged from 1.1 x 10^5^ to 3.2 x 10^7^ and 1.4 x 10^3^ to 1.6 x 10^7^, respectively, whereas *D. citri wg* gene copies ranged between 8.7 x 10^2^ to 2.4 x 10^3^ and 1.3 x 10^5^ to 1.2 x 10^6^, respectively. Thus, *D. citri* genomic DNA (which encodes the *wg* gene) was reduced by up to three orders of magnitude using the PDF protocol compared to the Direct protocol (Supplementary Fig. 2).

To assess the impact of host DNA removal on sequence coverage of *C*Las populations, we sequenced DNA samples prepared using the PDF and Direct protocols that possessed a range of *C*Las 16S rRNA gene copies for Illumina shotgun sequencing. The reduction in *D. citri wg* gene copies we detected also corresponded to a significant reduction in the average coverage of the *D. citri* genome (mean coverage of 0.3 and 0.9 for the PDF and Direct protocols, respectively; t_welch_ = 8.6, df = 33, p = 6.7 x 10^-10^) and a concomitant increase in sequencing effort afforded to *C*Las populations (Supplementary Fig. 2). In fact, when we compared the PDF protocol with existing protocols for sequencing *C*Las genomes available in the literature (Supplementary Table 1), the PDF protocol provided comparable coverage of *C*Las populations at orders of magnitude lower sequencing depth (Fig. 1a). Even for samples with high *C*Las titers (i.e., low Cq values observed for *C*Las 16S rRNA gene qPCR assays), average genome coverages of *C*Las populations in DNA samples generated using the PDF protocol were on average 35-fold greater (mean coverage of 43.2 and 1.24 for PDF and Direct protocols, respectively; t_welch_ = -3.37, df = 31, p = 0.002) than directly sequencing DNA extracted from an individual *D. citri* adult (“Direct protocol”, Fig. 1b). In addition to *C*Las, the PDF protocol significantly increased coverage of *Wolbachia* symbiont populations within *D. citri* adults by more than 7-fold compared to the Direct protocol (t_welch_ = -11.8, df = 33, p = 2.7 x 10^-13^) (Fig. 1b). While average coverage of *Wolbachia* symbiont populations approached 20X in the Direct protocol, they increased to 140X using the PDF protocol, greatly enhancing sequencing effort towards additional symbionts present within *D. citri* adults. Conversely, average coverage of the *D. citri* symbionts Car and Prof were significantly reduced by 6- and 3-fold, respectively, when comparing the PDF and Direct protocols, respectively. Nevertheless, average coverage of Car and Prof populations were 5X and 39X, respectively, using the PDF protocol which is sufficient for downstream genome assembly or comparative genomic analyses.

Aside from genome coverage estimation, we performed *de novo* genome assembly of individual and pooled samples (i.e., a coassembly) and examined the taxonomic identity of each contig by alignment to high quality reference genome sequences or NCBI’s nt database in concert with the BASTA tool for taxonomic annotation ^52^. Assembly statistics of *de novo* genome assemblies for all *D. citri* symbionts analyzed in the present study can be found in Supplemental Information (Supplementary Table 2). Of note, we assembled draft bacterial genomes from laboratory-reared *D. citri* adults assigned to the genus *Paracoccus* and *Brevundimonas* at 85 and 96% completeness, respectively, and with less than 5% contamination as assessed via the CheckM tool ^55^. Due to the uneven abundances of these taxa among *D. citri* individuals and their low coverage (Fig. 1b), their genomes were highly fragmented with low N50 values of 3,480 and 8,650 nt for *Paracoccus* and *Brevundimonas*, respectively, which are the sequence lengths at which sequences this long or longer compose at least 50% of the genome length. All assembled microbiome sequence data reported for laboratory-reared and field-collected *D. citri* adults (see below) can be accessed in the ‘Data availability’ section.

Next, we utilized the PDF protocol to sequence, assemble, and examine the microbiomes of field-collected *D. citri* adults (n = 6) collected from a citrus grove in Florida, USA in November, 2018. Unfortunately, *C*Las titers in these samples were quite low (*C*Las 16S rRNA gene Cq values > 30), and consequently any assembled *C*Las sequences detected in these samples were short in length and possessed low coverage (mean coverage ± s.d. = 0.74 ± 1.18). However, the coassembly of the six field-collected *D. citri* adults revealed large contigs assigned to the *D. citri* symbionts Car and Prof in each *D. citri* adult from the FL citrus grove (Supplementary Fig. 3a). Rather unexpectedly, we also detected 10,830 contigs (> 2,500 nt length) in the coassembly that were assigned to the *C. sinensis* genome, and in particular, large, high coverage contigs aligning to chloroplast and mitochondrial sequence elements of this same species (Supplementary Fig. 3b). Unlike our laboratory-reared specimens, in which we observed a significant sequence signal associated with the *D. citri* genome, only six contigs (>2,500 nt length) were assigned to the *D. citri* genome in field-collected adults, the longest of which was an ∼15 kb contig affiliated with the *D. citri* mitochondrial genome (Supplementary Fig. 3a). We also detected large numbers of low coverage contigs assigned to commensal bacterial populations including members of the genera, *Escherichia*, *Flavobacterium*, *Pseudomonas*, and *Xanthomonas*. Of note, only contigs assigned to the genus *Pseudomonas* achieved appreciably high coverage (∼20X) in one of the citrus grove-collected *D. citri* adults we sequenced (panel SPK8; Supplementary Fig. 3a). Finally, we observed high coverage contigs, some of which were large (∼15 – 30 kb), that were assigned to fungal mitochondrial sequences from well-known entomopathogenic fungal genera *Entomophthora*, *Capillidium*, *Neoconidiobolus* (Supplementary Fig. 3a). These sequences achieved appreciable coverage in only two of the six field-collected *D. citri* adults we sequenced (panels SPK1 and SPK4; Supplementary Fig. 3a), suggesting the PDF protocol may enable identification of *in situ* populations of entomopathogenic fungi potentially infecting *D. citri* in citrus groves.

In addition to shotgun sequence analysis and assembly of microbial genomes from lab-reared *D. citri* adults, we estimated replication rates for all four symbiont populations detected within individual *D. citri* sequenced using the PDF and Direct protocols. The genomic region predicted to contain the origin of replication (*oriC*) in bacteria can be distinguished by both GC skew (Fig. 2a, green line), a peak in average genome coverage (Fig. 2a, gray line), or conserved genetic elements associated with the *oriC*, usually *dnaA*, a gene involved in replication at the *oriC*, but whose genomic location differs in the alphaproteobacterial symbionts investigated herein and other loci are localized there instead (e.g., *hemE*, Fig. 2a) ^57,64^. Using the iRep tool, which performs regression analysis of ordered sequence read coverage over complete or draft microbial genome assemblies to estimate the index of replication (iRep), we were able to calculate an instantaneous measure of the average population replication rate ^57^ for the *D. citri* symbionts (Fig. 2b). The iRep values for the four symbionts were ordered Car > Prof > *C*Las > *Wolbachia* (Fig. 2c), but they differed between the PDF and Direct protocols. These differences were largely attributed to differential coverage between symbiont populations when comparing both protocols (Fig. 2d). For example, low sequence coverage of *C*Las and *Wolbachia* populations tended to result in higher iRep estimates in the Direct protocol data, whereas symbionts Car and Prof possessed higher coverage and lower iRep estimates with data produced by the Direct protocol. When comparing samples with coverage >20X, iRep values for Car and Prof populations in the PDF protocol data (1.52 ± 0.03 and 1.53 ± 0.11, respectively) approached an asymptote comparable to iRep estimates of Car and Prof populations in the Direct protocol data (1.67 ± 0.17 and 1.36 ± 0.09, respectively), but were not identical (Fig 2c, d). In contrast. *Wolbachia* populations achieved sufficient coverage (∼20X) for genome analysis in Direct protocol data, but their iRep estimates between the PDF and Direct protocol differed (1.33 ± 0.05 and 1.11 ± 0.02, respectively). To include samples from the Direct protocol, we chose to present unfiltered iRep values since coverage of some taxa, especially for the Direct protocol samples, failed to satisfy cutoff criteria required by the iRep tool to report an iRep estimate ^57^. For example, Car populations did not satisfy the iRep tool’s requirement that a minimum of 98% of genome windows possess 5X average coverage, even though the percentage of windows at this coverage were >93% in 39 out of 42 (93%) Car populations sequenced we sequenced. We also calculated iRep values for the handful of publicly available raw sequencing data available for *C*Las strains sequenced from *Citrus sinensis* and *Citrus maxima* DNA extracts, which ranged from 1.32 to 1.64 (Supplementary Table 3). However, average genome coverage of these *C*Las strains (see Methods for details) were between only 5 and 18X, which may inflate iRep estimates compared to *C*Las strains herein whose iRep estimates plateaued with increasing average genome coverage (Fig. 2d, PDF protocol panels). Overall, we observed that sequence data generated by the PDF protocol provides ample sequence coverage required to estimate stable iRep values for *C*Las (and other *D. citri* symbionts), especially when *C*Las titers in DNA extracts are sufficiently high (*C*Las 16S rRNA gene copies ≥1 x 10^5^) (Supplementary Fig. 2). All iRep estimates and associated metrics for symbionts in lab-reared *D. citri* and *C*Las genomes isolated from *Citrus* spp. can be found in Supplementary Table 3.

**Fig. 2.**
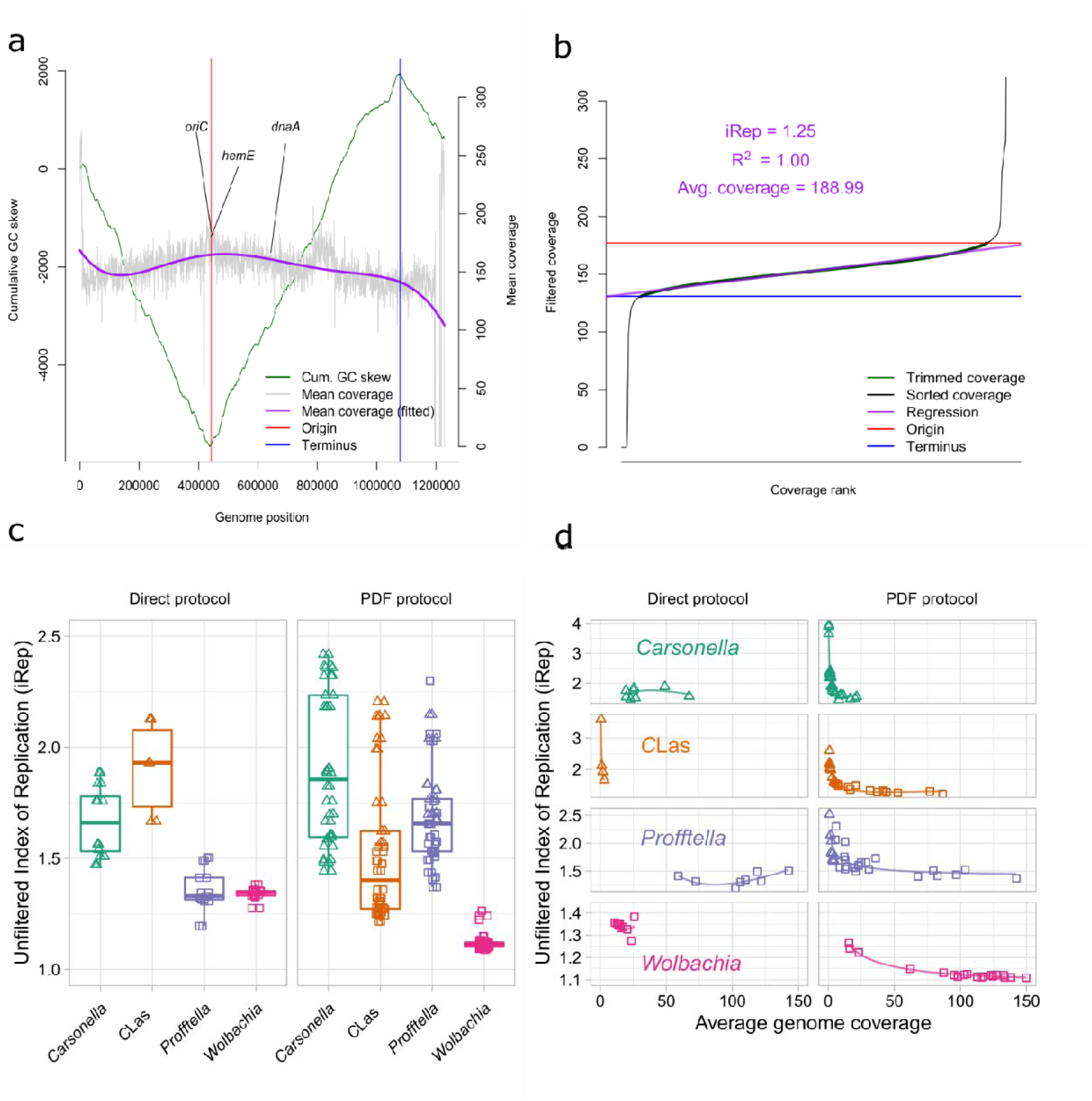
Index of replication (iRep) estimates for the four *D. citri* symbionts detected in sequence data sets generated from the PDF and Direct protocols. (a) The mean coverage (5 kb length windows, 10 nt sliding window) and GC skew across the *C*Las genome provide information about the replication origin (red line) and terminus (blue line). (b) The iRep tool ^57^ filters and sorts the coverage by rank along the *C*Las genome to derive a metric consistent with an average instantaneous replication rate within a bacterial population. (c) The iRep estimates were highest for ‘*Candidatus* Carsonella armatura’ in both the PDF and Direct protocols followed by ‘*Candidatus* Profftella armatura’, *C*Las, and *Wolbachia pipientis* strain wDi, but were shown to be highly dependent upon genome coverage (d). Squares and triangles in (c) and (d) indicate whether iRep could robustly calculate the filtered index of replication or not, respectively. Lines in (d) are second order polynomial regressions fit by formula iRep ∼ poly(log(coverage), 2) in R 4.1.2. Unfiltered index or replication values are shown to directly compare estimates from the PDF and Direct protocols. *oriC* = predicted origin or replication determined by the DoriC database ^97^ for *C*Las strain psy62; *hemE* = gene encoding uroporphyrinogen decarboxylase involved in heme biosynthesis and often detected adjacent to *oriC* in members of the *Alphaproteobacteria* ^64^; *dnaA* = encodes DnaA protein that binds to *dnaA* boxes to promote strand separation at the *oriC* region. All iRep data displayed in panels (c) and (d) be found in Supplementary Table 3.

Our next aim was to investigate the relationship of and genetic variation between *C*Las strains sequenced with the PDF protocol and publicly available *C*Las strains. Phylogenetic analysis of putatively non-recombinogenic SNPs identified with Gubbins ^59^ (see Methods for details) revealed a clear monophyletic signal among *C*Las strains sequenced with the PDF protocol (Fig. 3). The *C*Las strains sequenced from lab-reared *D. citri* adults in this study were observed to be descended from *C*Las strains (psy62 and JRPAMB1) derived from *D. citri* colonies maintained at USDA facilities in FL ^5^ (unpublished strain JRPAMB1 NCBI BioSample accession SAMN11842353; Fig. 3). In the 7 years since arriving in NY from FL rearing facilities, our *C*Las strains accumulated as many as 156 (78 ± 56, mean ± s.d.) genetic polymorphisms compared to reference *C*Las strain psy62 from FL (Fig. 4). In contrast, *C*Las sequence data produced using the Direct protocol in this study did not reveal detectable (min SNP coverage = 10) genetic polymorphisms in *C*Las, even in samples with high *C*Las titers (Fig. 1b, right panel). Of note, the PDF protocol enabled detection of genetic variants in two *C*Las genes predicted to be involved in antibiotic resistance by the ProGenomes database ^65^, a putative cysteine desulfurase and N-acetyltransferase (Fig. 4, purple text). *C*Las strains 20E and 22E possessed a missense mutation in the cysteine desulfurase *sufS* gene, at position 1,066,382, resulting in a base change of A to G and an associated amino acid sequence change of glutamine to arginine (amino acid 37 of *C*Las strain psy62 protein accession WP_015824969.1). We also detected a deletion mutation in the gene encoding a putative N-acetyltransferase (WP_102134467.1), in which a CA to C mutation at position 1,158,964 resulted in a frameshift mutation after amino acid position 36 in the encoded amino acid sequence. Interestingly, the gene (CLIBASIA_05345) encoding the putative N-acetyltransferase is located within a recently predicted prophage region of the *C*Las genome ^66^, and may constitute an important mobile element transferable among *C*Las strains if the prophage becomes lytic or is excised from the *C*Las genome ^67^.

**Fig. 3.**
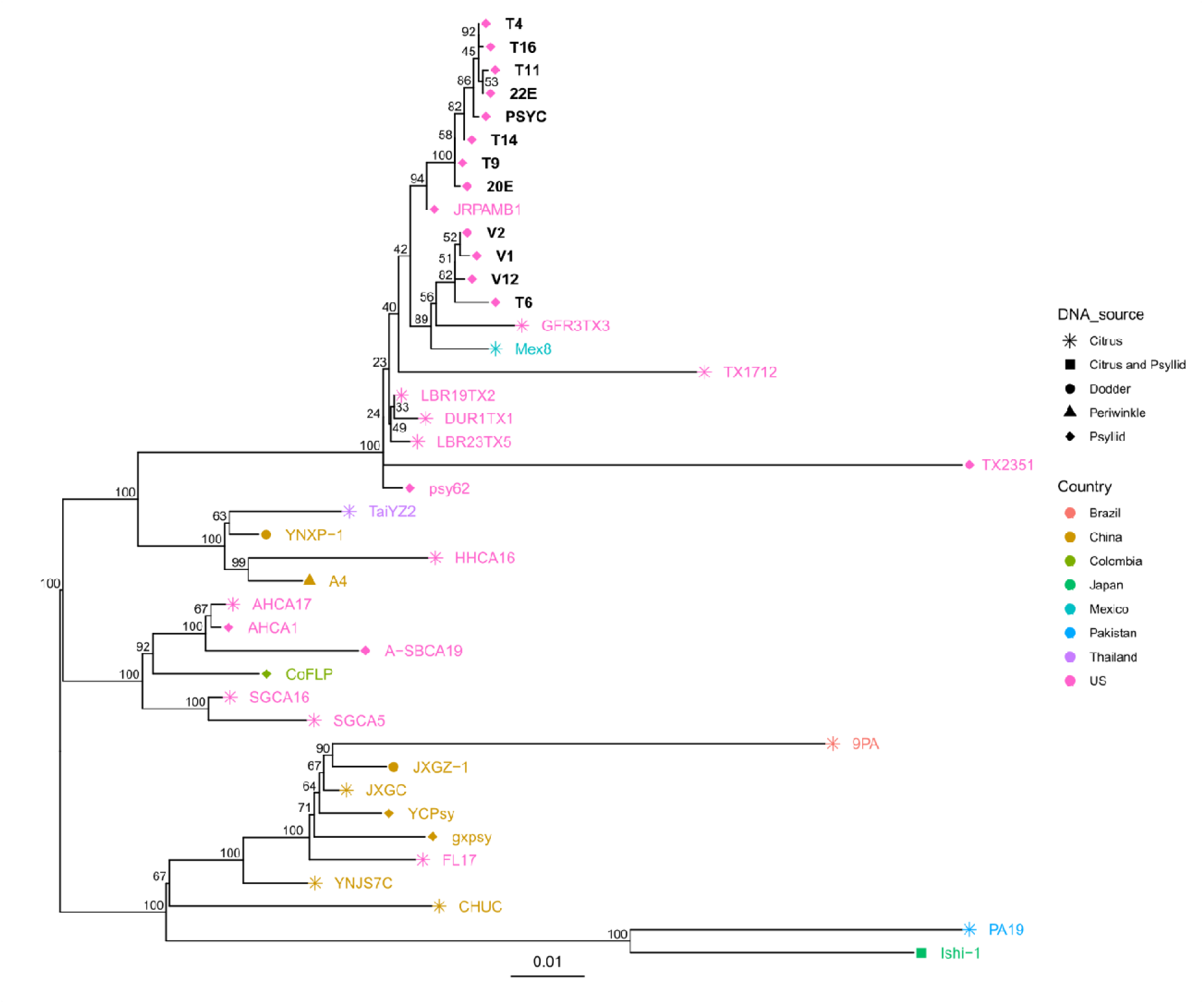
A midpoint-rooted, whole genome SNP phylogeny of the *C*Las strains derived from laboratory-reared *Diaphorina citri* adults using the PDF protocol (black tip labels) or *Citrus* spp. and *D. citri* tissues from public databases (colored tip labels). Shapes next to the tree tips indicate the DNA tissue source and colors of the shapes and tip labels represent the country of origin of the *C*Las genome (except black tip labels, which are US samples from this study). Values at nodes represent ultrafast bootstrap support reported by IQ-TREE ^60^ estimated using 1,000 bootstrap replicates. The scale bar represents the number of nucleotide substitutions per site. Strains with genome assemblies possessing median contig sizes of < 30 kb for more than half the contigs in the assembly were excluded, except for strain 9PA, which was included due to its unique geographic context in Brazil ^98^.

**Fig. 4.**
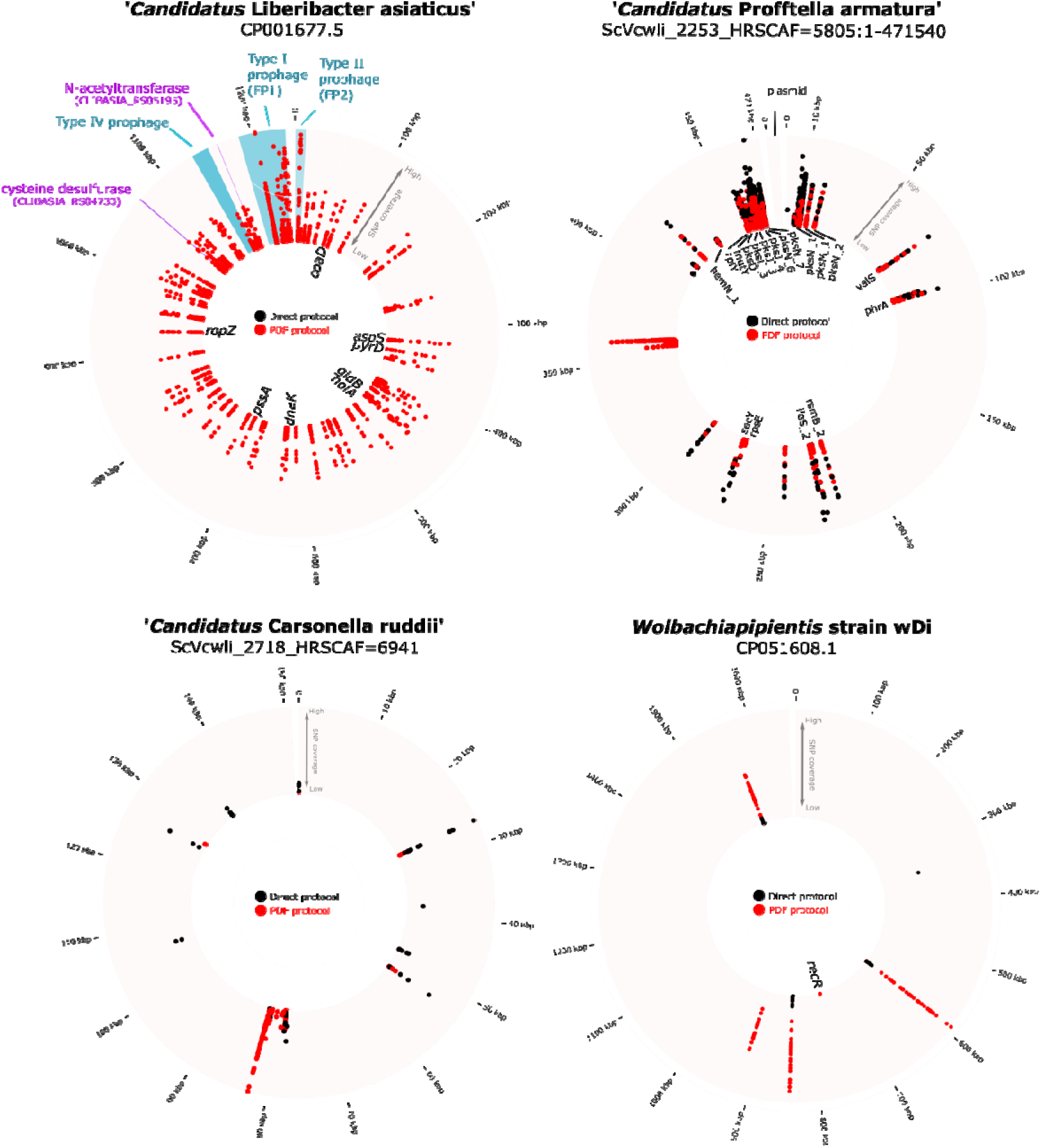
Coverage profiles of genetic polymorphisms detected within genomes of bacterial pathogens and symbionts of *Diaphorina citri*, the Asian citrus psyllid. Genetic polymorphisms identified in sequence reads generated using the PDF protocol (red points) versus the Direct protocol (black points) are compared. Teal-colored regions in the top left panel are predicted prophage-encoding regions of the *C*Las genome ^66^. Regions and labels in purple in the top left are two genes predicted to encode antibiotic resistance genes and observed to possess genetic polymorphisms compared to *C*Las strain psy62. Four letter codes inside each ring are gene abbreviations associated with annotated loci in which polymorphisms were detected. If a four letter abbreviation is absent, the polymorphism was either located within a gene lacking an annotation or was located in a noncoding region of the genome. All polymorphisms detected by snippy ^58^, including their four letter annotations, can be found in Supplementary Table 4.

In addition to potential antibiotic resistance, *C*Las encodes secreted protein effectors which may enable *C*Las to establish or persist during infection of its citrus or *D. citri* hosts ^34,68^. No variants were detected in loci encoding the secreted peroxidase SC2_gp095 (CLIBASIA_00485), the putative glutathione-dependent peroxidase SC2_gp100 (CLIBASIA_05595), or the Sec-translocon dependent effector SDE1 (CLIBASIA_05315). However, we did detect an insertion mutation in CLIBASIA_04530, encoding a putative Sec-dependent effector, which was shown to be predominantly expressed by *C*Las in citrus tissues ^34^ (Supplementary Table 4). While the exact functional consequences of the genetic polymorphisms detected between *C*Las strains in our lab and ancestral *C*Las strains in FL are unknown, the PDF protocol enabled robust detection and documentation of these mutations between *C*Las strains. Documentation of *C*Las genetic variants among laboratories should be a priority to ensure reproducible experimental outcomes among laboratories that maintain *C*Las strains via citrus grafting or acquisition and transmission of *C*Las by *D. citri*.

As previously noted*, C*Las strains also harbor distinct lytic and non-lytic prophage encoding regions which confer a variety of functions related to *C*Las pathogenicity and countering host immune responses ^33,66–68^. In the three predicted prophage elements FP1, FP2, and a recently discovered Type IV prophage element ^66^, we detected a total of 69, 11, and one polymorphisms in predicted coding regions, respectively, and only five polymorphisms in non-coding elements of the FP1 prophage region and none in the FP2 and the Type IV prophage element (Fig. 4). This was in contrast to non-prophage portions of the *C*Las genome, where only 33 polymorphisms were detected in coding regions compared to 52 in non-coding elements. The prophage variants occurred in phage primases, terminases, and four of the *C*Las strains we sequenced possessed an 8 bp insertion in the bacteriophage repressor protein c1, resulting in a frameshift mutation which could alter repression of the prophage lytic state within the strains this mutation was observed (Supplementary Table 4) ^69^.

Aside from *C*Las, the Prof symbiont was observed to possess the next greatest number of genetic polymorphisms when compared to the latest Prof reference genome from the citrusgreening.org database when using either protocol (Fig. 4). The location of genetic polymorphisms detected in Prof using either protocol generally agreed, although coverage was usually higher with the Direct protocol in Car and Prof populations and only a few unique polymorphisms were detected with each protocol (Fig. 4). Notably, a majority of the variants detected were frameshift or non-synonymous polymorphisms within genetic machinery involved in polyketide synthesis (Supplementary Table 4), which could indicate alteration to the defensive compounds (i.e., diaphorin) provided to *D. citri* by the Prof symbiont ^27^ and may be attributed to continuous laboratory propagation of *D. citri*. Genetic variants detected within Car, the nutritional symbiont of *D. citri* ^27,28^, were localized to genes encoding hypothetical proteins, RNA polymerase subunit beta’, DNA polymerase III subunit alpha, and the tRNA gene for phenylalanine (Supplementary Table 4). The genomes of *Wolbachia pipentis* strain wDi possessed the fewest number of polymorphisms (n = 6) compared to the *W. pipientis* strain wDi reference genome, with highest coverage of polymorphisms detected in samples generated using the PDF protocol (Fig. 4). However, all variants detected were frameshift or missense mutations in key genes putatively involved cell wall remodeling, conjugation, and cell regulation (Supplementary Table 4). We did not detect genetic variants in *Wolbachia* loci encoding proteins with a demonstrated interaction with *C*Las ^70^. All genetic variants detected in *D. citri* symbionts can be accessed in Supplementary Table 4.

## Discussion

Methods that enable or enhance sequencing of insect-associated microbiomes can inform solutions to longstanding challenges in diverse fields such as phytopathology, insect biology, and holobiont theory ^39,71^. For example, sequence-based analyses of microbiome composition and function can generate hypotheses related to the microbiome’s influence on insect fitness or the number and types of ecosystem services an insect and its microbiome (i.e., the holobiont) can provide ^39^. Furthermore, sequenced-based investigations of genetic modules underlying microbially-mediated pesticide resistance may facilitate new strategies to circumvent pesticide resistance, and may uncover novel resistance phenotypes of use to beneficial insects ^39,72^. Yet the high quantities of host DNA detected within insect microbiomes can hamper detection sensitivity of both microbial diversity and genetic polymorphisms present therein ^12^. Further, many insect-associated microorganisms of great importance are obligate or facultative symbionts that are as yet uncultivable ^73,74^, and therefore cultivation-independent methods to predict functions of uncultivable taxa are vital research tools ^35,71^. Thus, new approaches are needed to circumvent the limitations inherent to sequence-based analyses of insect microbiomes. The PDF protocol described herein enables cost-effective, efficient, and direct access to insect-associated microorganisms at orders of magnitude lower cost and total sequencing effort (Fig. 1a, Supplementary Table 1). Hence, the PDF protocol provides more microbiome data and facilitates investigations of microbe-microbe interactions or evolution at the molecular level.

Aside from the operational benefits afforded by the PDF protocol, it also enabled a comprehensive assessment of the microbiota present within and on *D. citri* adults. The PDF protocol not only enabled genetic analysis of both obligate (Car and Prof) and facultative (*C*Las and *Wolbachia*) symbionts, it also captured a variety of commensal bacterial and eukaryotic sequences in the *D. citri* adults examined (Fig. 1, Supplementary Fig. 3). The eukaryotic sequences we detected in field-collected *D. citri* are particularly intriguing as they represent genetic signatures of *Citrus* organelles retained due to *D. citri*’s phloem-feeding lifestyle. Thus, the shotgun sequence data provided by the PDF protocol may contain genetic signature’s consistent with an insect’s diet that could complement or supplant existing PCR-based approaches for gut content analysis in psyllids and other insects ^10^. The latter is particularly promising since PCR amplicons have limited value in taxonomic assignment due to their short length, whereas citrus chloroplast and mitochondrial contigs detected in PDF protocol data are tens to hundreds of kilobases in length, which will greatly assist with taxonomic identification of citrus or other plant species fed on by *D. citri*. Of note, stark differences in the composition of genetic signatures detected between the laboratory-reared and field-collected *D. citri* were observed. For example, laboratory-reared *D. citri* possessed much higher quantities of the *D. citri* genome than field-collected *D. citri* (Fig. 1b; except for a high coverage mitochondrial contig in field-collected *D. citri*, Supplementary Fig. 3a), whereas the field-collected *D. citri* data mostly contained contigs assigned to the *Citrus sinensis* genome (Supplementary Fig. 3b). The exact reasons for these differences are unknown, but they may be due to methodological differences in the capture and storage of laboratory-reared versus field-collected *D. citri*. For example, field-collected *D. citri* were preserved in 95% (v/v) ethanol when field-collected and then were subsequently frozen, whereas laboratory-reared *D. citri* were collected and frozen immediately. Thus, the cells of field-collected *D. citri* stored in ethanol prior to freezing may facilitate cell lysis, releasing more *D. citri* DNA that can be degraded during the DNase I treatment in the PDF protocol. The rigid wall of plant cells found in field-collected *D. citri* may be less affected by ethanol storage, and their cells were probably intact during DNase I treatment. This could explain why genetic signatures of citrus were higher in field-collected rather than laboratory-reared *D. citri*, if host cell lysis was less effective without ethanol storage or a freeze-thaw step. Nonetheless, these storage differences need to be explored in future iterations of the PDF protocol and may enhance digestion of *D. citri* or other insect DNA by DNase treatment.

In addition to genetic signatures indicative of the *D. citri* diet, we also detected mitochondrial signatures assigned to putatively entomopathogenic fungi, which suggests some members of the field-collected *D. citri* we studied were actively infected at the time of sampling (Supplementary Fig. 3). Interestingly, the mitochondrial signatures of the entomopathogenic fungi we detected are distinct from fungal genera previously reported to infect *D. citri* ^1^, and suggests additional, unexplored fungal taxa may be utilized as biocontrol agents for *D. citri* populations in the field. Fungi previously reported to infect *D. citri*, including members of the genera *Isaria*, *Hirsutellum*, *Lecanicillium*, and *Beauveria*, are all classified within the order Hypocreales, whereas the mitochondrial signatures we detected in the field-collected psyllids were all classified within the order Entomophthorales, members of which have been reported to infect *D. citri* in Mexico ^75,76^ but whose potential as biocontrol agents have not been explored. Despite the modest number of field-collected psyllids processed using the PDF protocol (n = 6), these abundant fungal signatures were also detectable, albeit at low coverage, within an additional 8 out of 17 (47%) *D. citri* adults that were collected from the same citrus grove, but were processed and sequenced using the Direct protocol (see Data availability to access raw data for the n = 17 *D. citri* individuals reported). These findings suggest an epizootic infection was occurring in the *D. citri* populations sampled from Spyke’s citrus grove in Florida at the time of collection. Therefore, the PDF protocol can facilitate the identification of native or potentially novel fungal pathogens of invasive insects, such as *D. citri*, and may assist the development of biocontrol strategies to prevent their spread ^77^.

Aside from *D. citri* microbiome composition, the sequence data afforded by our PDF protocol also enabled robust estimation of the replication rate of each symbiont present within the *D. citri* microbiome (Fig. 2). The same analysis is unlikely to be compatible with alternative bacterial enrichment protocols, such as multiple displacement amplification or probe capture hybridization methods. This is due to biased genome amplification and the variable capture efficiency among probes or loss of relative abundance among captured sequence elements for each method, respectively. However, probe hybridization is capable of enriching and sequencing *C*Las DNA from citrus tissues ^16^, which is currently unavailable with the PDF protocol described herein but is under active development. The latter two methods are the most widely used to sequence *C*Las genomes from *D. citri* adults or citrus samples, but both require greater time and financial investment and do not provide a comprehensive analysis of the *D. citri* microbiome. Nevertheless, the PDF protocol also has limitations, including reduced sequence coverage of obligate symbionts (Car and Prof) within *D. citri* compared to direct extraction and sequencing of DNA from *D. citri* adults using the Direct protocol. Despite this limitation, coverage of Car and Prof populations within *D. citri* adults enabled robust iRep estimates on par with those calculated from the Direct protocol (Fig. 2d). The estimation of *C*Las replication rates in *D. citri* adults is of interest to citrus greening disease research since actively replicating *C*Las populations are more likely to be spread by their *D. citri* vector ^78^. However, whether *C*Las actively replicates or only accumulates within its insect vector via feeding has been a matter of debate ^24,79^. According to iRep values recorded in the present study, approximately 20 to 30% of the *C*Las population with >25X coverage in the *D. citri* adults examined were actively replicating (Fig. 2). These values were much lower than Car and Prof populations, which at equivalent coverage (25X or greater) suggest that 50 to 75% of these symbiont populations are actively replicating, consistent with the key nutritional and defensive roles of these bacteria within *D. citri* ^26–28^. Considering the high prevalence of the *Wolbachia* symbiont within *D. citri* ^80,81^, we were surprised by the low iRep values estimated for this symbiont which suggests only 10% of the *Wolbachia* populations examined were actively replicating (Fig. 2). It’s possible that the unique mode of transovarial transmission by *Wolbachia*, competition with other symbionts, or inhibition by *D. citri*’s immune system might limit its replication within *D. citri* ^80,81^.

Despite its uncultivable nature, *C*Las is capable of propagation in laboratory settings using young citrus trees and laboratory-reared *D. citri* ^82–84^. While this indirect method of *C*Las cultivation has afforded valuable insights from therapeutics development to the response of *D. citri*’s immune system to *C*Las infection ^85–87^, continual propagation of bacteria in a laboratory can lead to bottleneck effects and introduce genetic variation not represented in founding or wild populations of the same bacteria ^88,89^. This fact is immediately apparent when comparing SNPs between reference *C*Las strain psy62 and the *C*Las strains sequenced in our laboratory (Fig. 3, 4). The great number of SNPs detected between *C*Las strain psy62 (a FL *C*Las strain) and our *C*Las strains in NY are concerning for several reasons. First, since our *C*Las strains derive from *C*Las strains maintained in FL greenhouses and *D. citri* rearing facilities, the number of differences suggest that our *C*Las strains have significantly diverged from strains in FL (average of ∼22 mutations/yr), which may result in inconsistent responses by therapeutics or other treatments being explored between facilities. To our knowledge, only a dozen *C*Las strains have been sequenced from commercial citrus groves or field-collected *D. citri* populations ^14,34, 90–92^. Rather, genome sequences of most *C*Las strains are derived from laboratory-reared *D. citri* infected with *C*Las or infected citrus scions grafted onto healthy citrus rootstock (Supplementary Table 1). This suggests our understanding of the existing genomic diversity of *C*Las strains in commercial groves or urban settings is poor. Furthermore, our understanding of genetic variation within and among laboratory *C*Las strains is vital to ensure reproducible results between institutions studying citrus greening disease and developing novel therapeutics to prevent *C*Las spread. Research conducted using institute-specific *C*Las strains may obfuscate or stymie investigations of *C*Las physiology or therapeutics development without dedicated methods to monitor and account for genetic variation among lab-propagated *C*Las strains. Our results and others ^93^ also support the idea that future studies take into account a role for *C*Las strain genetic diversity and propensity for mutation in fundamental studies on plant infection and psyllid transmission ^78^. Although hypothetical and difficult in practice, the development of approaches to inoculate plants and insects using standardized stocks of a diversity of *C*Las strains is warranted, as well as their storage and retrieval as viable stocks from a dedicated facility to promote reproducibility of results and accessibility of strains among institutions.

The biodiversity of insects is vastly underexplored and we have only yet to scratch the surface in our understanding of the relationship between insects and their microbial partners ^94^. Furthermore, there exists a paucity of methods to explore the host-microbe interface at the molecular level in insects and these methods are desperately needed for basic and applied entomological studies of insect vectors with medical and agricultural importance ^95,96^. We hope the PDF protocol and any future iterations thereof fulfills these needs and can enable advances in insect biology and microbiology to the benefit of all.

## Acknowledgements

SAH is grateful to Dr. Ashley M. Frank for assistance with revising the manuscript. SAH would also like to thank Dr. Hannah J. MacLeod and Stephanie Preising for their helpful scientific discussions regarding the development of the PDF protocol. All authors are grateful to Dr. Stacy DeBlasio, Dr. David Igwe, Luke Thompson, and Julie Blaha for their tireless efforts maintaining and provisioning the citrus colonies and *C*Las-infected *D. citri* adults utilized in this work. This research used resources provided by the SCINet project of the USDA Agricultural Research Service, ARS project number 0500-00093-001-00-D. We gratefully acknowledge the financial support of the California Citrus Research Board grant # 5300-196, the USDA ARS Administrator’s Class of 2021 Research Associate Program, a USDA NIFA predoctoral fellowship to MM and the USDA Agricultural Research Service CRIS Project 8062-22410-007-000-D for funding.

