## Supplemental Information for "Direct DNA sequencing of ‘*Candidatus* Liberibacter asiaticus’ from *Diaphorina citri*, the Asian citrus psyllid, and its implications for citrus greening disease management"

**Supplementary Figures**

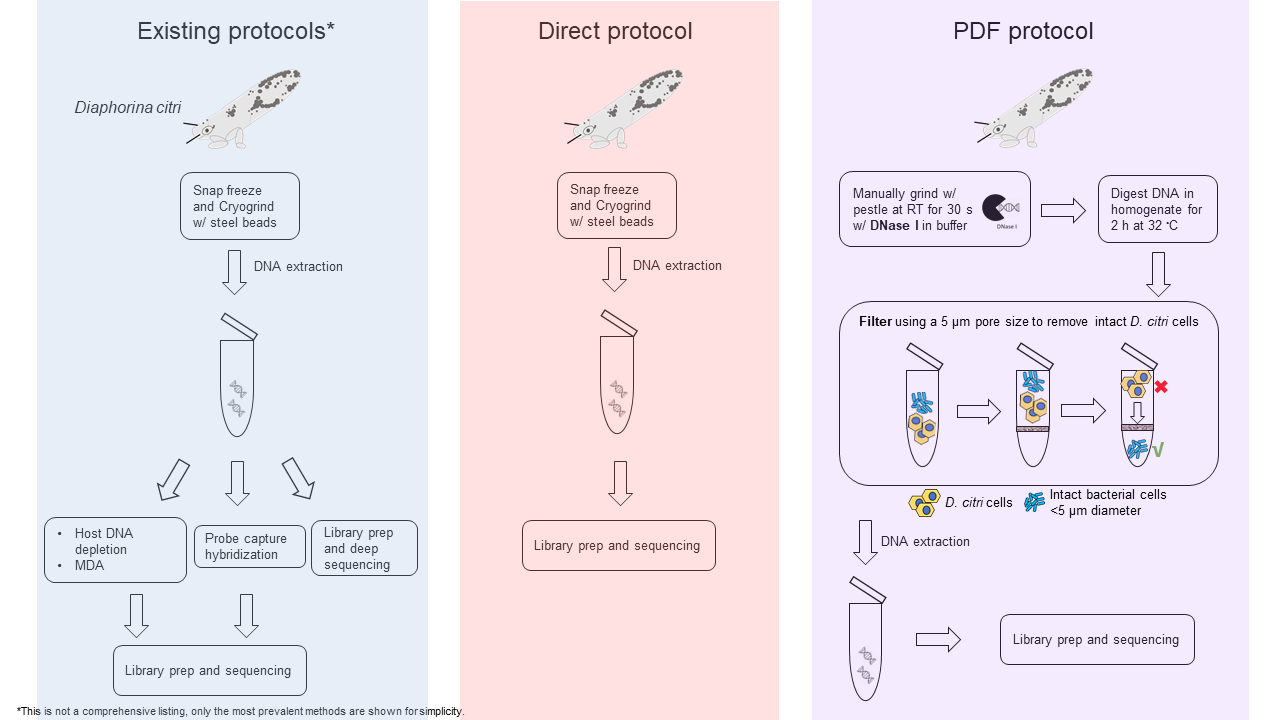

**Supplementary Fig. 1.** Overview of the protocols to enrich and sequence ‘*Candidatus* Liberibacter asiaticus’ (*C*Las) from its insect host, the Asian citrus psyllid, *Diaphorina citri*. The existing protocols to enrich and sequence *C*Las in the literature include post DNA extraction treatments, including host DNA depletion by methylation status (host DNA depletion, above) followed by multiple displacement amplification (MDA), use of complementary RNA probes targeting the *C*Las genome via probe capture hybridization, or deep sequencing of *Citrus* spp. or *D. citri* tissues with high *C*Las titers. The Direct protocol includes directly extracting DNA from *C*Las-infected *D. citri* and submitting for library preparation and DNA sequencing. The Direct protocol is used for comparison to the pre-treatment DNase and filtration (PDF) protocol in the main text. Finally, the PDF protocol includes manually grinding *D. citri* tissues using a microcentrifuge pestle in a buffer containing DNase I enzyme, digestion of DNA for 2 h, following by filtration of the cell lysate through a five micron pore size filter by centrifugation. Then, DNA extraction is performed on the resulting cell pellet, followed by library preparation and sequencing. The asterisk next to the “Existing protocols” in the figure indicates that this is not a comprehensive listing of all methods to enrich and sequence *C*Las from its hosts, only that the most prevalent approaches have been presented for simplicity.

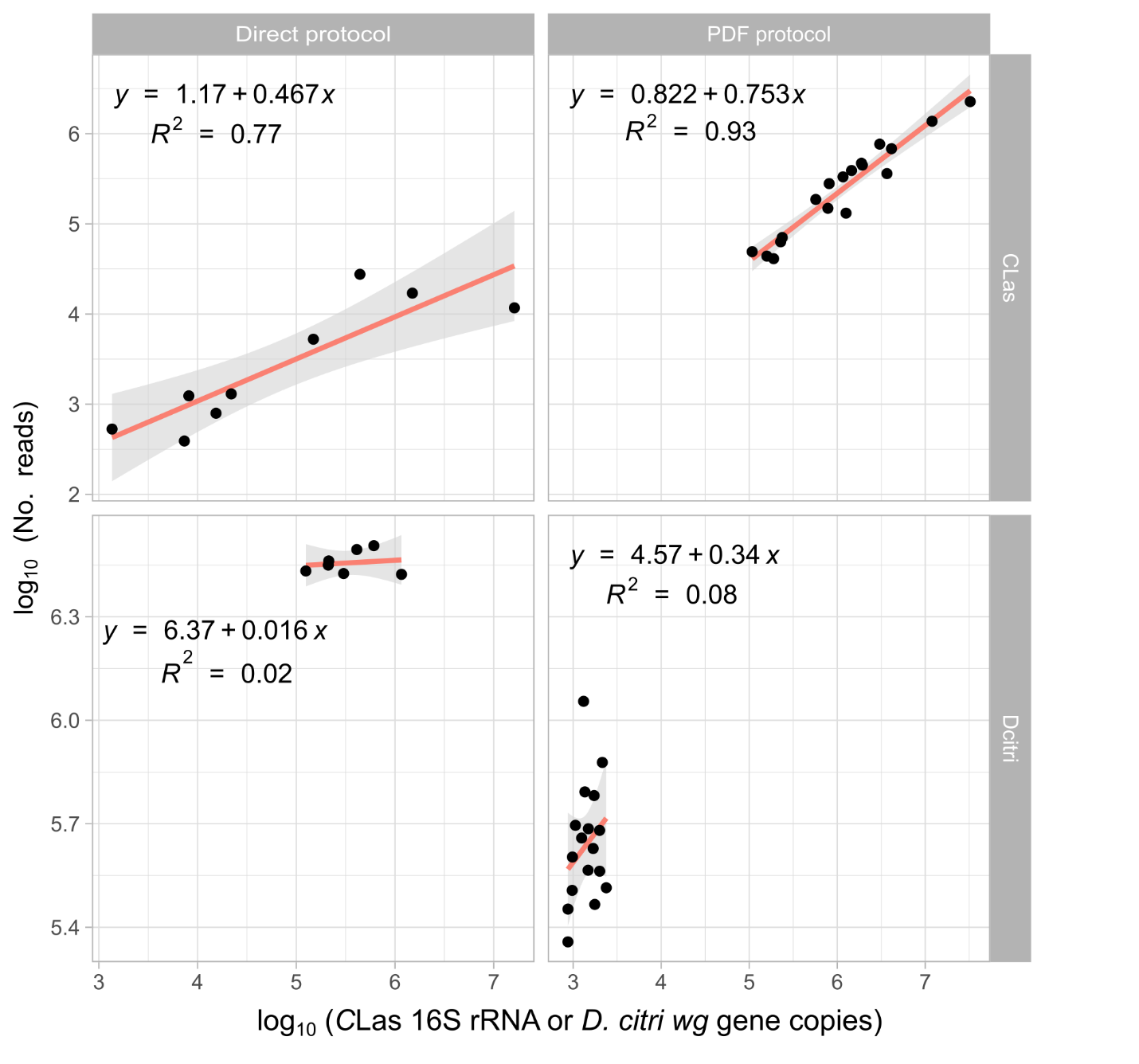

**Supplementary Fig. 2.** Relationship between the number of Illumina reads aligning to ‘*Candidatus* Liberibacter asiaticus’ strain psy62 (*C*Las) or the *Diaphorina citri* v3 genome and *C*Las 16S rRNA or *D. citri wg* (*wingless*) gene copies. The terms “PDF protocol” and “Direct protocol” above each plot refer to whether individual psyllid samples were filtered (5 micron pore size) and treated with DNase I prior to DNA extraction or whether DNA was directly extracted from *D. citri* individuals without pre-processing (see main text Methods for details). *C*Las = “*Candidatus* Liberibacter asiaticus” and Dcitri = *Diaphorina citri*. The red line and grey shading surrounding the line in each plot represents the linear regression fit and 95% confidence interval estimates, respectively. The linear regression equation and goodness-of-fit statistic is presented in black text within each panel. Note the differences in scale on the y-axis panel labels.

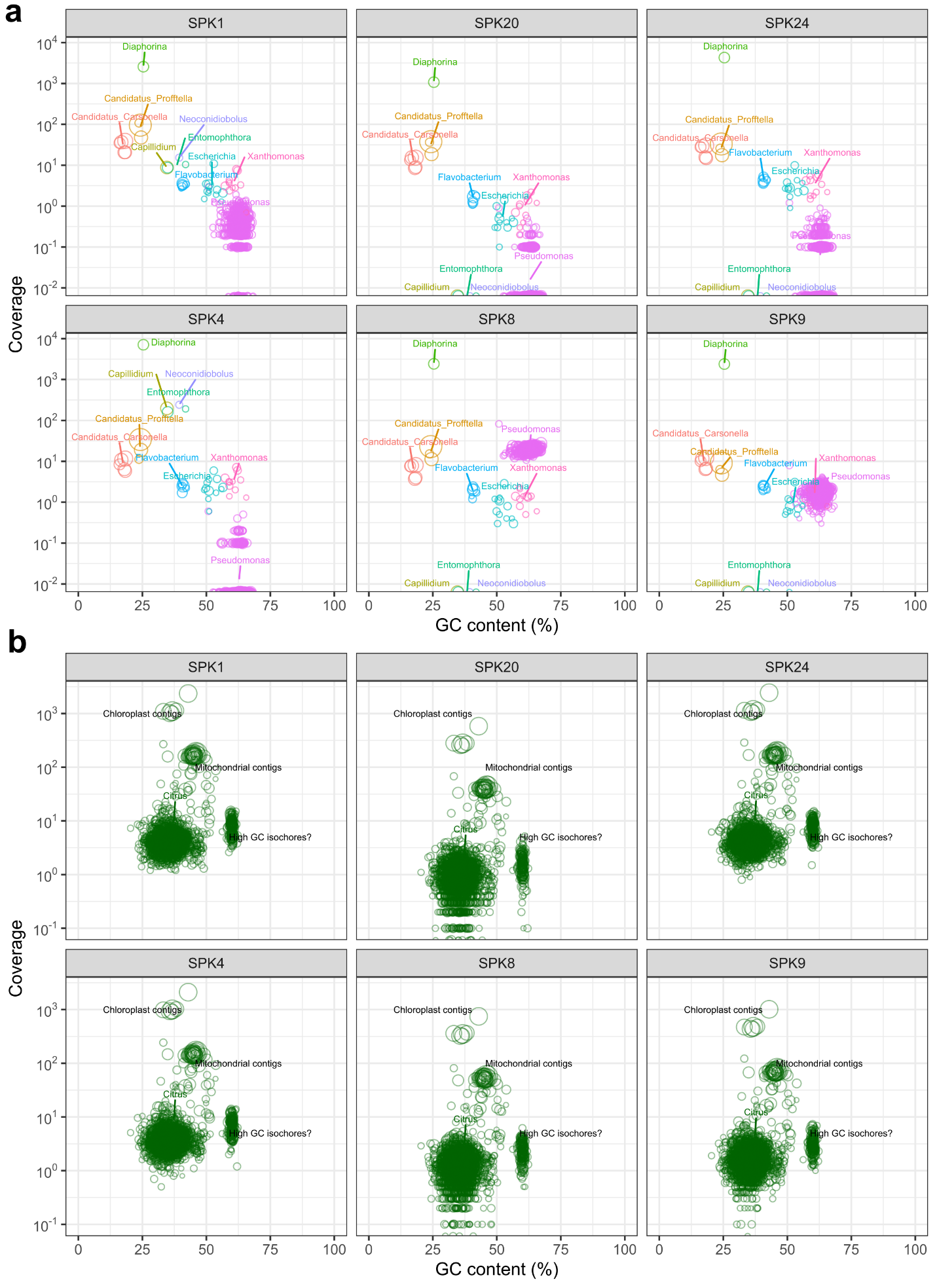

**Supplementary Fig. 3.** Taxonomic composition, coverage, size, and % GC content of assembled contigs (>2,500 nt) from six individual field-collect *D. citri* adults processed using the PDF protocol. The size and color of points in all panels represent the length and taxonomic affiliation of contigs generated from a coassembly of sequence reads from all individuals. The assembled contigs were taxonomically assigned to *D. citri* (mitochondrial contigs), entomopathogenic fungi (*Capillidium*, *Neoconidiobolus*, *Entomophthora*), and *D. citri* symbionts or commensal bacterial taxa (a) or to the genomes of *Citrus* *sinensis* or *C. maxima* (b). Note the variation in coverage of contigs assigned to putative entomopathogenic fungi or the bacterial genus *Pseudomonas* in (a) and the consistently high coverage of citrus chloroplast and mitochondrial contigs in (b).

**Supplementary Tables**

See Extended Data Excel files for the following tables.

**Supplementary Table 1.** Sequencing effort and cost required to sequence *C*Las from *C*Las-infected insect (*Diaphorina citri*) or plant (*Citrus* spp.) tissues.

**Supplementary Table 2.** Assembly statistics for *Diaphorina citri* symbionts assembled using the SPAdes assembler.

**Supplementary Table 3.** Index of replication (iRep) estimates and associated metrics from lab-reared *Diaphorina citri* symbionts and publicly available *C*Las genomes from *Citrus* spp.

**Supplementary Table 4.** Genetic polymorphisms detected in symbionts of *Diaphorina citri* adults.
